# Confocal Raman Microscopy-Guided Optimization of Early Otic Differentiation from Human Pluripotent Stem Cells

**DOI:** 10.64898/2026.06.24.734338

**Authors:** Keshi Chung, Damien Veret, Duong Phu Le, Ludivine Rouillon, Elias Estephan, Alban Desoutter, Hamideh Salehi, Azel Zine

**Affiliations:** LBN, Laboratory of Bioengineering and Nanoscience, Univ Montpellier, 34000 Montpellier, France; Cube, UMR 7357, CNRS, INSERM, University of Strasbourg, 67000 Strasbourg, France

## Abstract

Generation of otic progenitors from pluripotent stem cells requires precise timed regulation of signalling pathways, including bone morphogenetic protein 4 (BMP4). Because endogenous levels of BMP4 varie between cell lines, the optimal concentration of exogenous BMP4 must be determined individually to achieve efficient otic differentiation. Three different human induced pluripotent stem cell lines (hiPSCs) underwent ectodermal differentiation to early otic induction stages in the presence of various concentrations of BMP4 (0-5 ng/ml). Differentiation outcomes were assessed by immunofluorescence staining, and quantitative gene expression analysis. Raman microscopy was used to characterize biochemical differences between hiPSC differentiated cultures exposed to different BMP4 concentration. We observed distinct ectodermal fate were after 8 days of in vitro differentiation depending on BMP4 concentration, including neural, non-neural/otic ectoderm and surface epidermal fates. The proportion of PAX2-otic progenitors varied substantially between cell lines and culture conditions, ranging from approximately 9% to 77%. Raman spectroscopy revealed concentration dependent spectral differences and enabled discrimination between differentiating condition within individual hiPSC lines. Analysis of Raman spectral features indicated differences in nucleic acid, lipid, protein, and collagen associated signatures across culture conditions and cell lines. These findings demonstrate that Raman microscopy provides a non-destructive, label-free method for monitoring molecular changes associated with early otic differentiation. By complementing conventional molecular and immunocytochemical analyses, Raman spectroscopy offers a valuable tool for optimizing BMP4-mediated otic induction protocols and improving the reproducibility of stem cell-based strategies for inner ear research and regenerative medicine.

## Introduction

Sensorineural hearing loss is one of the most common sensory disorders worldwide and is primarily caused by the degeneration of cochlear hair cells, spiral ganglion neurons, or other specialised cell types in the inner ear which do not regenerate.^1,2^ As a result, attempts have been made to generate otic organoids from human pluripotent stem cells to produce otic cells for transplantation, to better understand human otic development, and as a model for screening compounds for potential otoprotective and/or otoregenerative effects aimed at restoring auditory function.

Human pluripotent stem cells (hPSCs), including human induced pluripotent stem cells (hiPSCs), provide a unique platform for generating inner-ear cell types in vitro.

Over the past decade, significant advances have been made in directing hPSCs toward otic lineages, the generation of otic progenitors, sensory hair-cell-like cells, and spiral ganglion neuron-like cells in otic organoids.^3–12^ Despite these advances, the otic differentiation procedures remain time-consuming and difficult to replicate across hPSC lines.^13,14^ This is compounded by heterogeneity as a result of differences in different hPSC being used both within and between laboratories. Indeed, because of different levels of endogenous signalling molecules, such as BMP4, between hPSCs, differentiation must be optimised for each cell line that is used in a study.^15^ This is particularly crucial during the early stages of otic differentiation, when levels of BMP4 that are too low drive differentiation towards neural fate, while high levels of BMP4 drive differentiation towards surface epidermal fate. It is therefore necessary that the concentration of BMP4 is optimised for each cell line to achieve robust and reproducible otic differentiation outcomes. Current methods of determining the optimal concentration of BMP4 for otic induction typically involve immunocytochemistry and gene expression analysis of otic organoids at early stages of differentiation for the presence of target (non-neural ectoderm) and off-target (neural ectoderm and surface epiderm) markers.^3^ Screening of early otic organoids for epidermal thickness has also been used to optimise BMP4 concentration.^12^ However, these methods they are often labour-intensive, destructive, and require the generation of large numbers of organoids in several differentiating conditions, the consumption of large amounts of culture media and antibodies.

Given these limitations, there is a need for methods that can rapidly and non-invasively assess differentiation status without consuming large quantities of reagents or requiring extensive sample processing. Raman microscopy fulfils these criteria: it is label-free, compatible with both live and fixed cells, and has been successfully applied to characterise stem cells undergoing differentiation towards various lineages^16–20^, as well as to monitor critical quality attributes during cell therapy manufacturing.^21^

Here we propose its use an alternative and complementary approach for optimising BMP4 concentrations during early otic induction. Beyond its practical utility, Raman imaging offers the additional advantage of revealing changes in the biochemical composition of cells as they acquire distinct ectodermal identities, thereby contributing to a deeper understanding of the molecular events underlying early otic specification in the human inner ear.

## Results

### Induction of ectodermal fates

The three human iPSC lines used in this study for ectodermal induction and otic cell differentiation (ASE, CHIPS, and SCT) (**Figure 1 A-C**) were first confirmed to have different endogenous levels of BMP4 expression using qPCR analyses (**Figure 2B**). Prior to induction of ectodermal differentiation, all cell lines were validated for the expression of pluripotency markers. Following 8 days of differentiation in 2D cultures, expression of pluripotency markers *OCT4* and *NANOG* had decreased in all cell lines at all concentrations of BMP4 tested (**Figure 2A, Figure S1**), indicating a loss of pluripotency following the induction of differentiation towards ectodermal fates.

**Figure 1.**
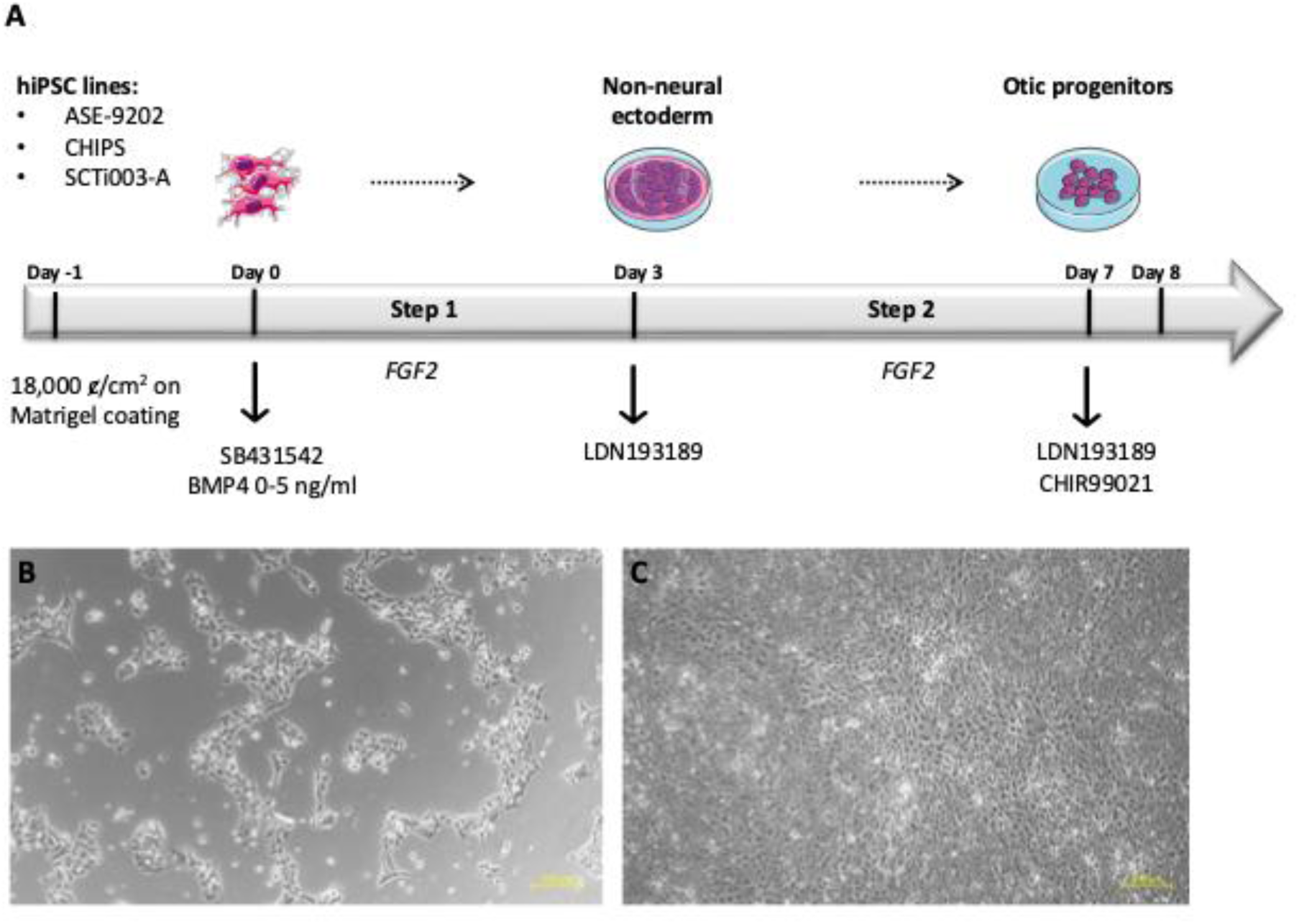
Schematic summary outlining the otic differentiation from hiPSCs in 2D cell cultures. (**A**) As a first step (step1), hiPSCs at day 0 were exposed to SB431542 (TGFb inhibition) and varying concentrations of BMP4 (0-5 ng/ml) until day 3 for early no-neural ectoderm induction and, in a second step (step2) were then differentiated into otic progenitor cells by exposure to both LDN (BMP4 inhibition) and CHIR99021 (WNT activation) until day 8. (**B**-**C**) Representative phase contrast images for morphological characteristics of hiPSC-derived otic progenitor cells. A: propagated hiPSCs (ASC line) at day 0 before differentiation. B: differentiated otic cells at day 8 in BMP4 (1 ng/ml) treated cultures. Scale bars: 200 µm.

**Figure 2.**
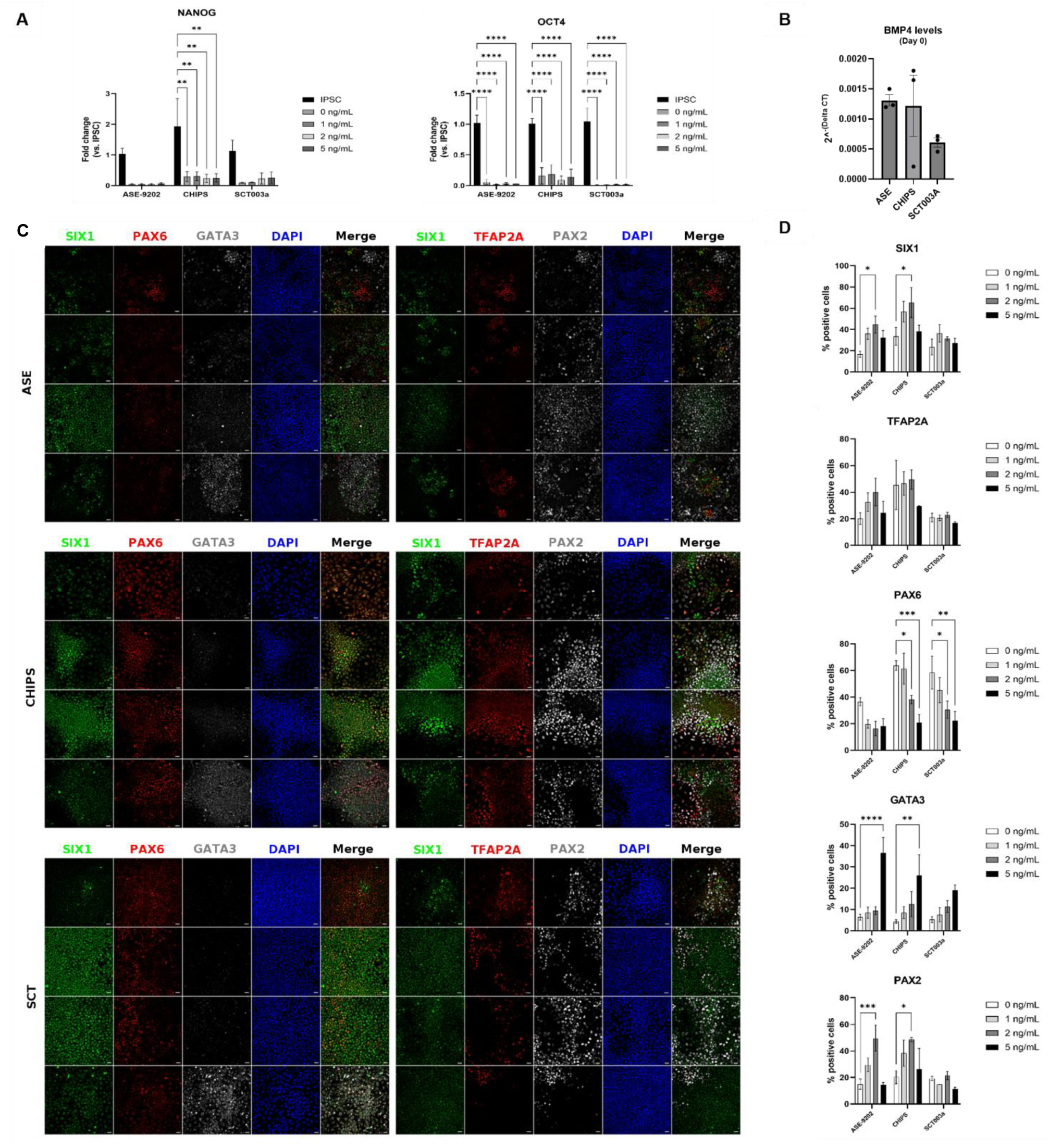
Induction of ectodermal fates in iPSCs. (**A**) Loss of pluripotent identity after 8 days as indicated by reduced expression of OCT4 and NANOG mRNA compared to GAPDH as housekeeping gene (n = 3 biological replicates) (**B**) Estimation of the BMP4 levels at the iPSC states (day 0) for each cell lines compared to GAPDH as housekeeping gene (**C**) Images of cells stained for PAX6 neuronal ectoderm marker, GATA3 surface ectoderm, SIX1 and TFAP2A non-neural ectoderm, and PAX2-otic progenitors (**D**) and their related quantifications (n = 3 to 5 biological replicates). Statistical analyses were performed by 2-way ANOVA for panel A and D and by Kruskal-Wallis test for panel B. * p-value ≤ 0.05, ** p-value ≤0.01, *** p-value ≤0.001, **** p-value ≤ 0.0001.

Expression of the neural ectodermal marker PAX6 was highest in all cell lines exposed to 0 ng/ml BMP4 during the early stages of differentiation (**Figure 2**). Expression of PAX6 decreased as higher concentrations of BMP4 were used, suggesting a loss of neuronal identity. In contrast, GATA3 expression was highest in all cell lines when 5 ng/ml BMP4 was used, indicative of differentiation towards surface epidermal fate, which was not observed in the presence of lower concentrations of BMP4. Intermediate concentrations of BMP4 tended to result in increased expression of the non-neural ectodermal markers SIX1 and TFAP2A, as well as the otic progenitor marker PAX2 (**Figure 2 C-D**). This effect was particularly pronounced for the ASE and CHIPS lines. Exposure to 1 ng/ml to 2 ng/ml BMP4 therefore appeared to be optimal for induction of otic differentiation for all cell lines used in the study.

### Raman analysis of differentiated hiPSC lines

While immunocytochemical methods can be used to determine the optimal concentration of BMP4 for otic induction, we wondered whether an alternative method might be used that could potentially be faster and more informative about changes in the biochemical properties of the cells as they differentiate. We therefore performed Raman imaging on the cell cultures of all 3 cell lines 8 days after in vitro differentiation using the same protocol as above. PCA was used to investigate the relationship between them and different concentrations of BMP4.

### ASE cell line

Principal component analysis (PCA) revealed concentration-dependent spectral changes associated with BMP4 treatment (**Figure 3A-C**). The first principal component (PC1), accounting for 17.44% of the total variance, showed a tendency to separate the 0 ng/ml group from the 1 and 5 ng/ml BMP4-treated groups. However, given the relatively low proportion of explained variance, this separation should be interpreted cautiously. Notably, samples exposed to higher BMP4 concentrations exhibited tighter clustering than the untreated group, suggesting reduced spectral variability and the emergence of a more homogeneous biological response. Comparison of the mean spectra and PCA loading plots identified several Raman bands contributing to group discrimination, including peaks at 727, 793, 920–963, 1008, 1304, 1347, 1439, 1680, 2849, 2893–2905, and 2945 cm⁻¹.

**Figure 3.**
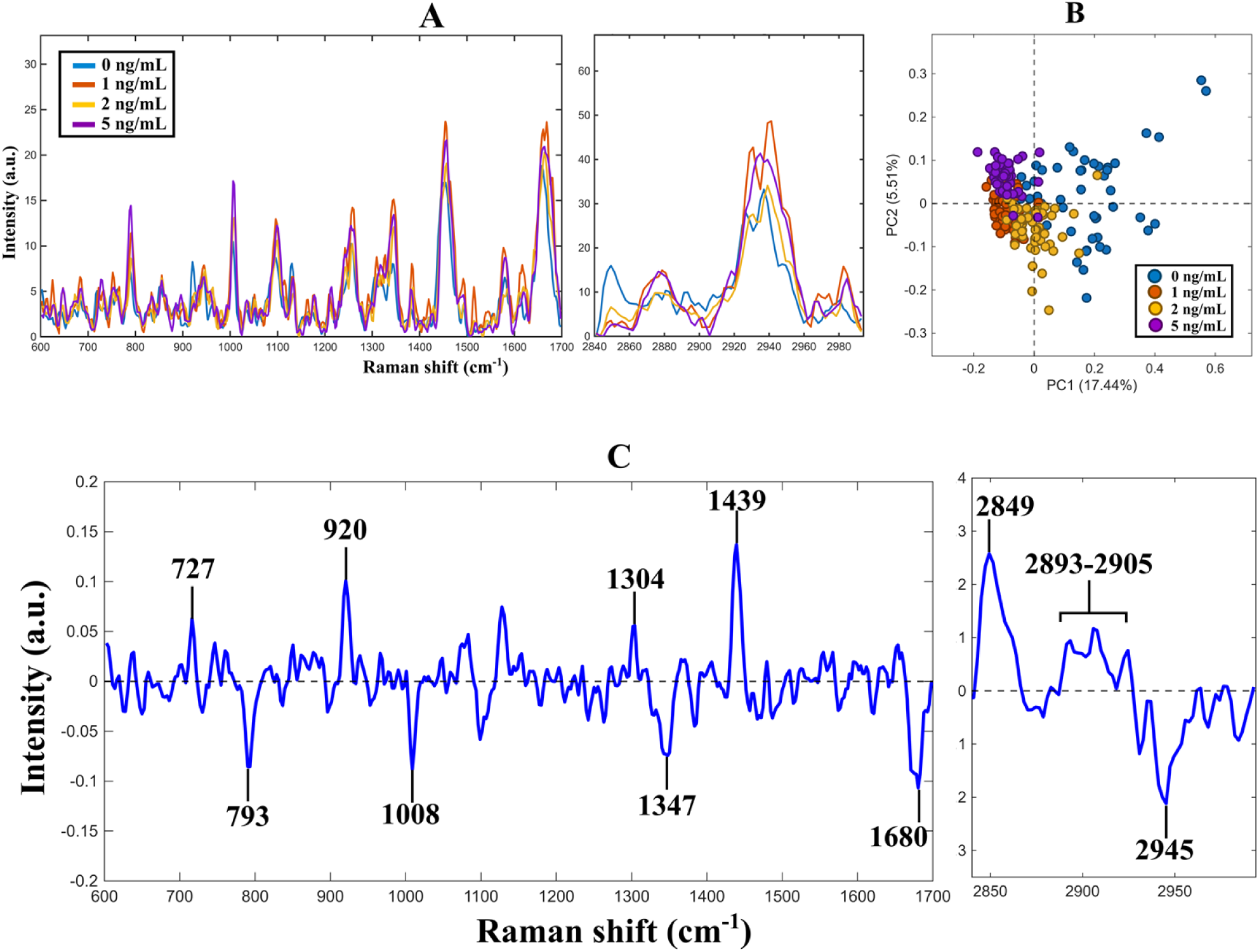
Analysis in ASE line cell: (**A**) mean spectra, (**B**) score plot of PCA result, (**C**) loading of PC1.

To further visualize biochemical and structural changes during differentiation, false-colored Raman images were generated using the Proline band (854 cm⁻¹) and the C-H stretching band (∼2940 cm⁻¹), the latter being mainly associated with proteins and lipids. Merged images of these signals revealed distinct spatial organizations and structural features that occurred during cell differentiation (**Figure S2**).

To obtain a more robust characterization of the spectral changes, a feature-based approach was adopted. Rather than focusing on individual Raman bands, this strategy integrates multiple spectral features and relies on peak intensity ratios, thereby reducing the influence of noise and normalization-related bias. Four major biochemical feature groups were analyzed: nucleic acids, proteins, lipids, and collagen, with collagen considered separately because of its specific role in extracellular matrix organization. The boxplot analysis (Figure 4) revealed distinct BMP4 concentration-dependent trends among these biochemical features. Signals associated with nucleic acids progressively increased with BMP4 concentration, reaching their highest values at 5 ng/ml. In contrast, protein- and lipid-related features displayed opposing trends. Protein-associated signals generally increased with increasing BMP4 concentration, although a transient decrease was observed at 2 ng/ml. Conversely, lipid-associated features showed an overall decline, most pronounced at 5 ng/ml. Collagen-related features followed a pattern similar to that observed for proteins, exhibiting an overall increase with BMP4 concentration while also showing a reduction at 2 ng/ml.

**Figure 4.**
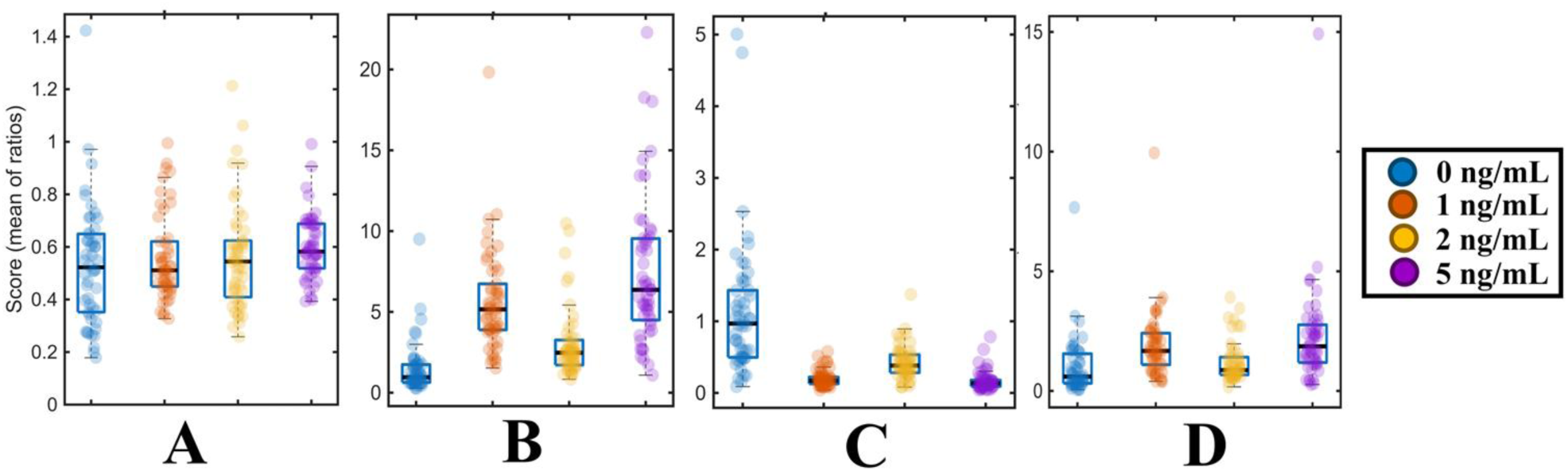
Ratio scores: associated-DNA (**A**), associated-protein (**B**), associated-lipid (**C**), associated-collagen **(D**) in the ASE cell line.

### CHIPS cell line

For the CHIPS cell line, the PCA results (**Figure 5B**) indicate that the first principal component (PC1, explaining 8.19% of the variance) shows a tendency to separate the 2 ng/ml treatment group from the remaining groups. However, given the relatively low explained variance, this separation should be interpreted as partial. The second principal component (PC2, 5.86%) also suggests some degree of differentiation between the 0 ng/ml and 5 ng/ml groups, although the separation is less pronounced compared to PC1. Analysis of the loading plots in conjunction with the mean spectra indicates that the observed differences are primarily associated with four main spectral feature regions: nucleic acids, proteins, lipids, and collagen.

**Figure 5.**
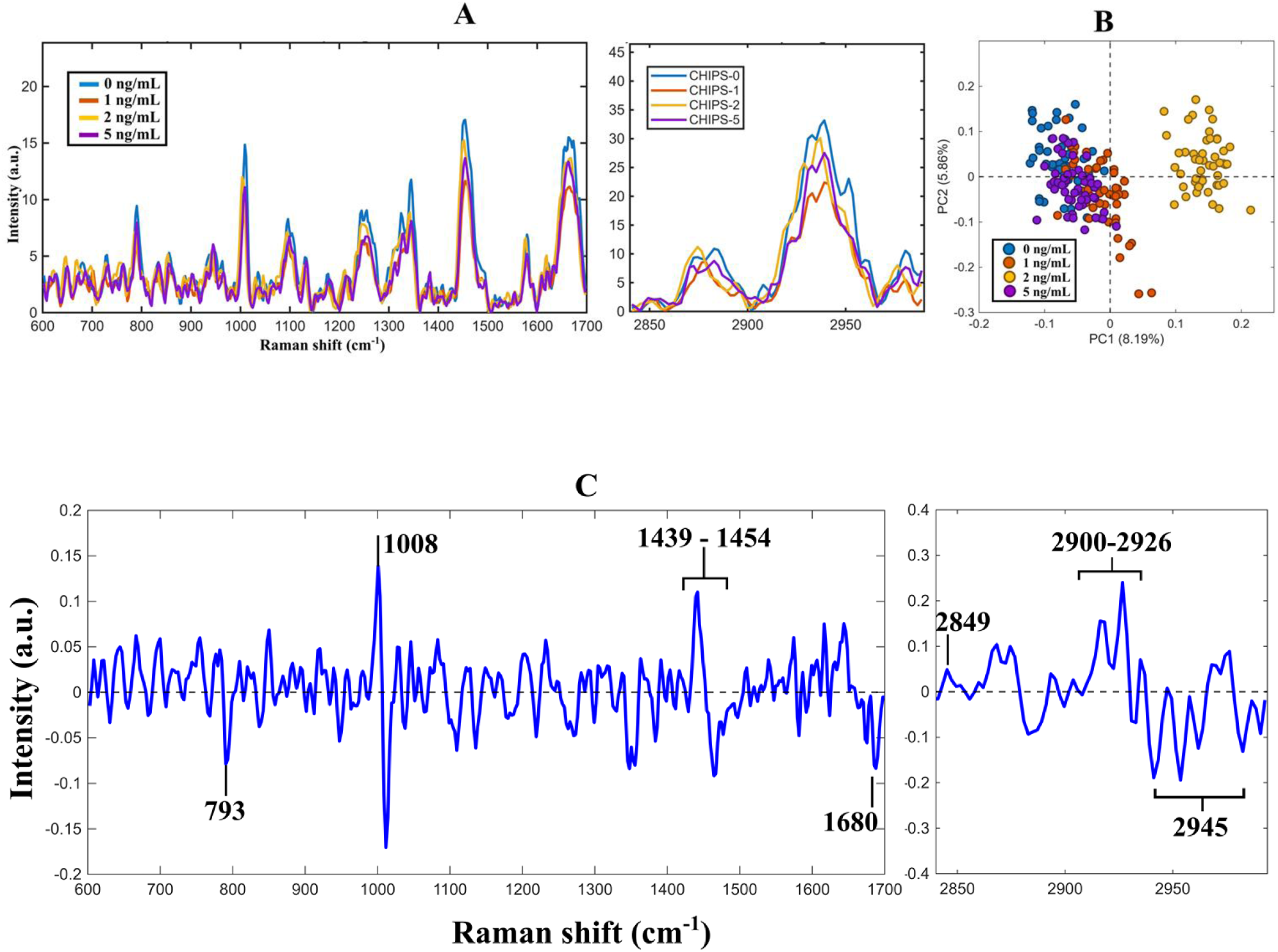
Analysis in CHIPS line cell: (**A**) mean spectra, (**B**) score plot of PCA result, (**C**) loading of PC1.

The results in **Figure 6** indicate a slight increasing trend in the nucleic acid-related region with increasing BMP4 concentration; however, this trend is less pronounced compared to that observed in the ASE cell line. The protein- and lipid-related regions continue to exhibit opposing trends with respect to BMP4 concentration, with protein-associated signals tending to decrease and lipid-associated signals increasing at higher concentrations. The collagen-related region shows a pattern consistent with that of protein. Overall, the magnitude of differences among groups in the CHIPS cell line appears less pronounced than in the ASE cell line, suggesting that the biological response to BMP4 may differ in strength or sensitivity between the two hiPSC lines.

**Figure 6.**
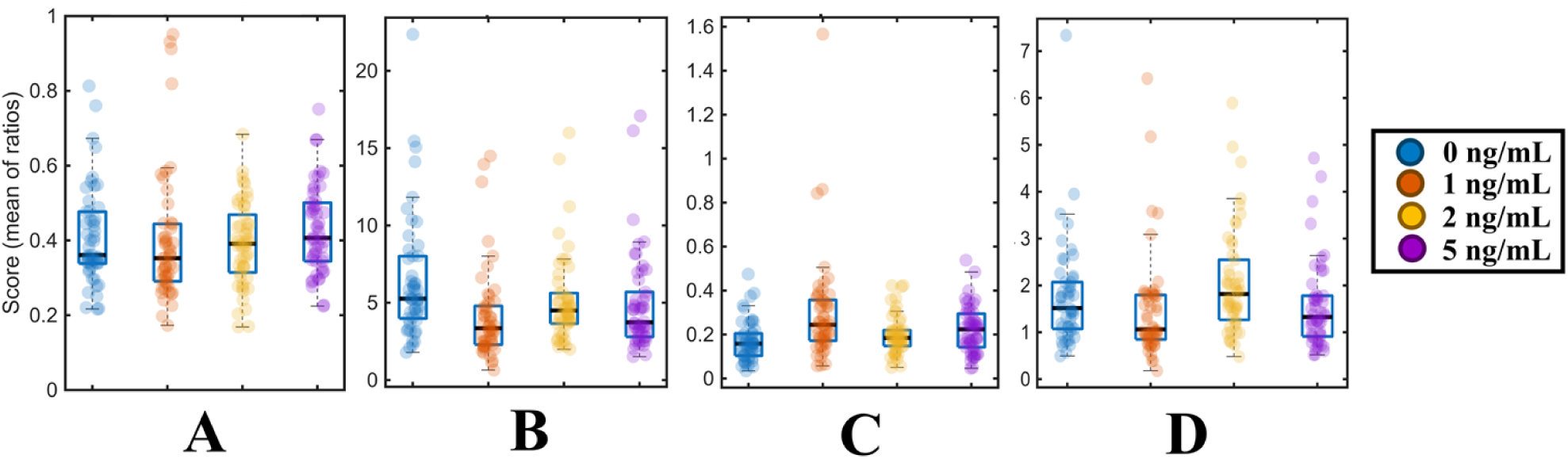
Ratio scores: associated-DNA (**A**), associated-protein (**B**), associated-lipid (**C**), associated-collagen (**D**) in the CHIPS cell line.

### SCT cell line

The results for the SCT cell line are presented in **Figure 7 and Figure 8**. PCA analysis (**Figure 7B**) indicates that the fourth principal component (PC4, explaining 1.9% of the variance) suggests only a very limited separation between the 0 ng/ml and 5 ng/ml groups. Given the low explained variance, the discriminative power is weak. The distribution of data points shows a greater degree of overlap among clusters, making them less distinguishable compared to the ASE and CHIPS cell lines, which suggests lower variability between treatment conditions. Comparison of the PC4 loading profiles with the mean spectra indicates that spectral regions associated with nucleic acids, proteins, lipids, and collagen still contribute to the loadings, although the overall differences between groups remain subtle. In the SCT cell line (**Figure 8**), the nucleic acid-related region shows a more pronounced increasing trend compared to that observed in the CHIPS cell line, with values rising as BMP4 concentration increases. The protein-related region exhibits a decreasing trend with increasing BMP4 concentration, whereas the lipid-related region shows an opposite pattern, increasing with higher BMP4 levels. Consistent with the previous two cell lines, the collagen-related region in the SCT cell line follows a trend similar to that of the protein-related region, with values decreasing as BMP4 concentration increases.

**Figure 7.**
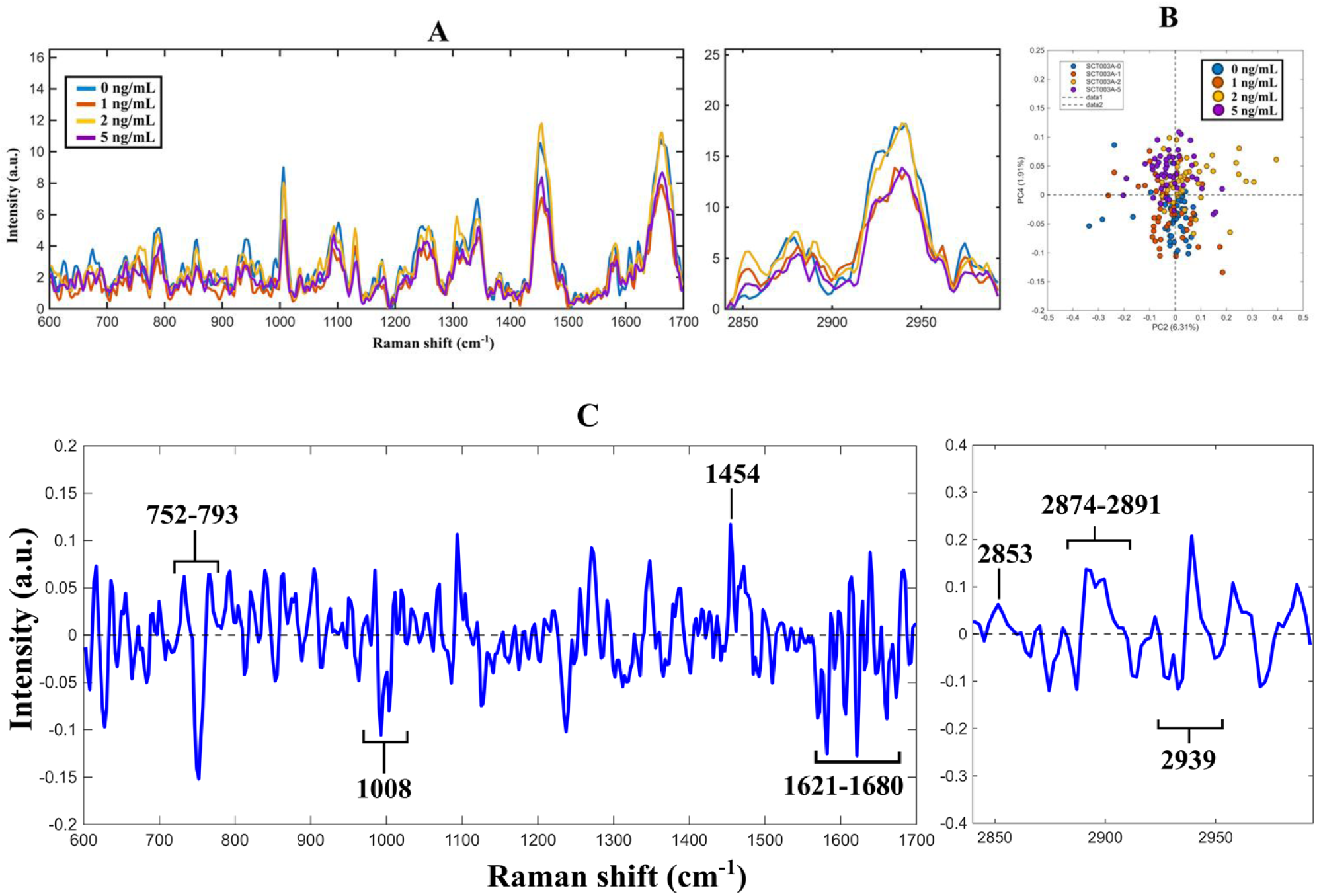
Analysis in SCT line cell: (**A**) mean spectra, (**B**) score plot of PCA result, (**C**) loading of PC1

**Figure 8.**
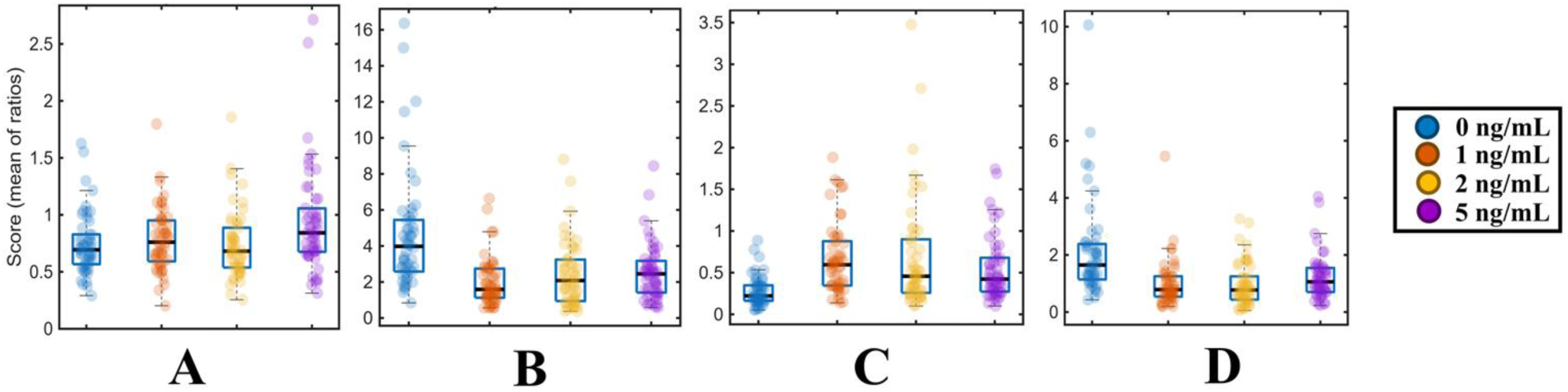
Ratio scores: associated-DNA (**A**), associated-protein (**B**), associated-lipid (**C**), associated-collagen (**D**) in the SCT cell line

### Comparison of Raman signatures across differentiated hiPSC lines

Comparison of Raman-derived indices across all three hiPSC lines revealed both shared and cell-line-specific responses to BMP4 exposure. The most consistent feature was a progressive increase in nucleic acid-associated signals with increasing BMP4 concentration, which was observed in ASE, CHIPS and SCT cultures.

In contrast, protein, lipid and collagen-associated signatures differed between cell lines. The ASE line exhibited increasing protein and collagen-associated indices together with decreasing lipid-associated indices at higher BMP4 concentrations. Conversely, the CHIPS and SCT lines displayed decreasing protein- and collagen-associated indices accompanied by increasing lipid-associated signals. These findings demonstrate that although BMP4 exposure induces common biochemical changes across hiPSC lines, substantial cell-line-specific variation remains evident in the Raman spectral response.

## Discussion

In this study, we demonstrate that Raman microscopy can distinguish between neural, non-neural, and surface epidermal ectodermal fates induced in hiPSCs by varying concentrations of BMP4, and that this discrimination is reflected in characteristic spectral signatures associated with DNA, protein, lipid, and collagen content. These findings position Raman imaging as a practical and informative complement to existing methods for optimising otic induction protocols. A key challenge in the field is that differentiation outcomes are highly dependent on the endogenous signalling environment of each cell line in particular, on baseline BMP4 levels meaning that exogenous BMP4 concentrations must be individually calibrated. Previous studies have shown that differing levels of endogenous signalling pathways, including BMP4 and WNT, affects the amounts of exogenous small molecules that needs to be added to each cell line in order for differentiation to proceed as intended.^3,12,22^

We demonstrate that different hiPSC lines exhibit distinct responses to exogenous BMP4 during early ectodermal differentiation, resulting in variable efficiencies of otic induction. We further show that Raman microscopy detects reproducible biochemical differences associated with these differentiation outcomes monitoring early ectodermal fate specification and optimising differentiation conditions in a cell-line-specific manner.

Consistent with previous reports, differentiation outcomes were strongly influenced by BMP4 concentration. Low BMP4 levels favoured neural ectodermal differentiation, whereas high BMP4 concentrations promoted epidermal fate, while intermediate concentrations resulted in the greatest expression of non-neural ectodermal and otic markers. These observations further support the concept that graded BMP signalling governs ectodermal fate decisions and highlight the importance of optimising BMP4 exposure for individual cell lines. The differences observed among the ASE, CHIPS and SCT lines are likely attributable, at least in part, to variations in endogenous signalling activity, which have previously been reported to influence lineage specification in pluripotent stem-cell cultures. Such variability remains a major challenge for the reproducibility of differentiation protocols and reinforces the need for robust methods capable of rapidly identifying optimal culture conditions. Among the spectral features identified, we observed that DNA, lipid, and protein (including collagen) signatures varied depending on the cell line and specific ectodermal fate that was being induced, and that it could be correlated with cell fate.

Specifically, across all three cell lines, consistent nucleic acid-associated signatures tended to be low with lower concentrations of BMP4 (corresponding to neuronal induction) and increased with increasing BMP4 concentration (corresponding to surface epiderm induction). Non-neuronal (presumptive otic) induction fate had intermediate levels of nucleic acid contributions, which was between that of neuronal and surface epidermal fates. This could reflect changes in transcriptional activity as a result of induction of specific ectodermal fates.

Several biological mechanisms may contribute to these observations, including changes in chromatin organisation, transcriptional activity, DNA/RNA content, or cell-cycle dynamics.

For instance, the chromatin remodeller CHD7, which is already expressed at the non-neural ectoderm stage of otic induction, is known to be important for regulating key genes associated with otic development.^23^ Transcription factors such as VAX2, POU3F2, FOXN2 and ZBTB18 are important for neuronal differentiation.^24^ Differences in the expression and activity of chromatin remodellers and transcription factors during the induction of different cell fates would result in differences in nuclear content and potentially unique signatures that could be used as readouts of specific cell fates. Understanding how these differences in DNA signatures are related to different cell fates could help in elucidating the mechanisms of induction and differentiation at an individual cellular level of ectodermal lineage specification.

The protein, lipid, and collagen-associated signatures exhibited more complex behaviour and difference between cell lines. A low protein and collagen index was associated with neuronal induction for the ASE cell line, while higher protein and collagen indices were indicative of non-neuronal and surface epiderm induction. The opposite was observed for the CHIPS and SCT lines, where higher protein and collagen indices corresponded to neuronal induction. The reason for this discrepancy is not clear, but as the differentiation protocol was optimised for non-neural/early otic differentiation over the other cell fates, incomplete and/or impartial neural induction could mean that not all cells in the culture had fully acquired a neuronal cell fate at the same time across the different cell lines.

Interestingly, the ASE line showed increasing protein and collagen-associated signals with increasing BMP4 concentration, whereas the CHIPS and SCT lines displayed the opposite trend. This divergence likely reflects intrinsic differences in cellular responses between hiPSC lines and emphasises the heterogeneous nature of pluripotent stem-cell differentiation. Importantly, these differences further support the rationale for cell-line-specific optimisation strategies. Rather than identifying universal biochemical markers of differentiation, Raman analysis may provide a means of defining line-specific spectral signatures associated with successful otic induction.

The collagen-associated spectral features observed in this study are of particular interest given the established role of extracellular matrix ECM) components during inner-ear development. Previous studies have demonstrated the expression of multiple collagen isoforms in the rodent.^25,26^ and human fetal cochlea.^27^ More recently, mutations affecting collagen genes (i.e. COL2A1) have been linked to hearing impairment.^28^ Expression of type-II collagen is absent in the tectorial membrane of mice with disrupted expression of *COL9A1*, suggesting that interaction of different collagens interact with each other is essential for proper development and functioning of the inner ear. Indeed, type-II collagen in particular has been found to be an important constituent of the extracellular matrix (ECM) of the developing inner ear of mice.^29^

Interestingly, in a previous transcriptome study, we reported dynamic expression changes of several of ECM, including collagens coincident with hiPSC differentiation towards human otic neurosensory fates.^30^ It is therefore conceivable that collagens play an essential role during early otic induction and are required at optimal concentrations to ensure an appropriate environment for proper inner ear development.

The present findings suggest that the ECM remodeling may accompany early ectodermal fate specification. However, since collagen-related assignments were inferred from Raman spectral features rather than direct molecular measurements, further studies will be required to identify the specific ECM components contributing to these signals and to determine their functional significance during otic development.

Similarly, the lipid-associated signatures detected by Raman microscopy may reflect important metabolic and structural changes occurring during differentiation. The Raman imaging data also suggests that the lipid signal is important and characteristic during induction of different cell fates, although to a lesser extent than that of protein and collagen.

Lipids are abundantly expressed in different cell types and their levels must be appropriately regulated for proper cell differentiation, maturation, function, and survival. Intracellular lipid storage organelles called lipid droplets are found in neurogenic niches, where they regulate neural stem cell proliferation and are dynamically remodelled upon fate changes such as quiescence and differentiation^31^ and have been observed to have an antioxidant role during hypoxia and oxidative stress during development in Drosophila.^32^

In contrast, hyperactive neurons have been reported to transfer toxic fatty acids to astrocytes, protecting neurons during periods of enhanced activity.^33^ This suggests that lipids are potentially both protective and harmful in neurons and requiring their levels to be optimally regulated. Whether such regulation of lipids is also essential in the inner ear remains to be investigated, although we speculate that this may be the case given the presence and importance of lipids in this system.

Interestingly, recent evidence has also implicated lipid-mediated regulation of FGF/MAPK signalling pathways during otic placode formation.^34^ Additionally, lipid-rich microdomains in both sensory and non-sensory cells in the inner ear are known to be essential for cochlear function^35^, emphasising the importance of lipids in the inner ear.

The differential lipid signatures observed across differentiation conditions therefore suggest that lipid metabolism may contribute to lineage specification during early otic development. Future studies integrating Raman imaging with lipidomic and transcriptomic analyses could provide further insight into these mechanisms.

Because Raman imaging is label-free, it is suitable for use in cells that are intended for further downstream applications and potentially even for clinical use. The cells that were used in this study were fixed prior to imaging. Future studies should aim to replicate the finding in living cells that can then be returned to culture conditions for further differentiation and ensure that early optimisation of induction of non-neural ectoderm translates to improved otic induction and differentiation of hair cells in otic organoids.

Although Raman spectroscopy clearly detected biochemical differences between differentiation conditions, future work should evaluate the predictive performance of Raman-based classification approaches using larger datasets and supervised machine-learning methods.

In conclusion, this study showed that Raman microscopy is a powerful, non-invasive technique that can distinguish between different induced ectodermal fates of stem cells. It also confirms that each cell line behaves differently during induction to different ectodermal fates based on exposure to different BMP4 concentrations, complementing observations made using other techniques. Moreover, the technique offers additional insight into the characteristics and mechanisms of cells as they differentiate towards their specific fates, making it a promising technique for advancing the stem cell field and regenerative medicine.

## Materials and Methods

### Validation of human induced pluripotent stem cell lines

The commercially sourced cell lines used in this study were ASE-9202 (ASE) (Applied Stem Cell), ChiPSC-4 (CHIPS) (Cellartis, Takara Bio Europe), and SCTi003A (SCT) (Stem Cell Technologies). The ASE-9202 line was generated from human skin fibroblasts using non-integrating episomal vectors. ChiPSC-4 was reprogrammed from foreskin fibroblasts of a healthy human donor by using retrovirus technology based on the transcription factors Oct3/4, Sox2, Klf4, and c-Myc.^36,37^ The SCTi003A line was derived from peripheral blood mononuclear cells from a healthy female donor.

Cell lines ASE and CHIPS were karyotyped by Stem Genomics (Montpellier, France) prior to performing experiments (**Figure S3**). The SCT line was recently acquired from the supplier at low passage and was not karyotyped at the time of conducting the experiments. All lines were validated for pluripotency markers using immunofluorescence staining for SOX2, OCT4, and SSEA4 using the Plurtipotent Stem Cell 4-Marker. Immunohistochemistry Kit (Invitrogen) (**Figure S1**), as well as qPCR for *OCT4*, *NANOG*, and *SOX2*, and were regularly tested to ensure that they were free from mycoplasma contamination using MycoStrips (Invivogen).

### Human induced pluripotent stem cell cultures

The hiPSC lines ASE, CHIPS, and SCT at passages less than 30 were cultured on tissue culture dishes coated with Matrigel (Corning) at approximately 8 μm/cm^2^ in mTeSR Plus medium (Stem Cell Technologies). Cells were maintained in a humidified incubator at 37°C with 5% CO_2_. Culture medium was changed every second day, and cells were passaged when they reached about 70% confluency (approximately every 5-7 days) using 0.5 mM EDTA (Sigma).

### Ectodermal differentiation

Differentiation of hiPSC lines to neural ectoderm, non-neural ectoderm/early otic fate, and surface epiderm was achieved using a modified protocol for ectodermal differentiation towards otic fate.^3,5,7,12^ Briefly, colonies were dissociated into single cells using Accutase (Thermo Fisher Scientific) and seeded at a density of 18,000 cells per cm^2^ on to either glass coverslips (for immunofluorescence or qPCR analysis) or CaF_2_ coverslips (for Raman imaging) coated with Matrigel in mTeSR Plus supplemented with 10 µM Y27632 ROCK inhibitor (MedChemExpress) (day -2). When cells had reached about 60% confluency (usually 48 hours after seeding), cells were washed once with E6 medium and then cultured for 3 days in E6 medium supplemented with 4 ng/ml FGF2 (Stem Cell Technologies), 10 μM SB431542 (Stemgent), 100 μg/ml Normocin (Invivogen), and BMP4 (Reprocell) at concentrations ranging from 0 ng/ml to 5 ng/ml (day 0). On day 3 of differentiation, the medium was replaced with E6 medium supplemented with 50 ng/ml FGF2, 200 nM LDN193189 (Stemgent), and 100 μg/ml Normocin. On day 7, the differentiation medium was replaced with E6 medium supplemented with 3 μM CHIR99021 (Reprocell), 50 ng/ml FGF2, 200 nM LDN193189, and 100 μg/ml Normocin. Cells were collected on day 8 for downstream applications.

### Immunofluorescence and microscopy

Cells were washed in PBS and fixed in 4% PFA for 10 minutes at room temperature. Permeabilisation was performed with 0.3% Triton X-100 (Sigma) in PBS for 1 hour at room temperature, followed by blocking in 10% donkey serum for 1 hour at room temperature. Apart from the SOX2, OCT4, and SSEA4 antibodies provided with the Pluripotent Stem Cell 4-Marker Immunohistochemistry Kit (Invitrogen) which were prepared according to the manufacturer’s protocol, primary antibodies (**Table S1**) were diluted in blocking buffer, and cells were subsequently incubated in primary antibody dilutions overnight at 4°C. Cells were then washed 3 times for 5 minutes with 0.3% Triton X-100 in PBS, followed by incubation in appropriate secondary antibodies diluted in blocking buffer for 2 hours at room temperature. This was followed by 10-minute incubation in 1 μg/ml DAPI (Sigma) and three 5 minutes washed with 0.3% Triton X-100 in PBS. Finally, cells were mounted on glass slides in Fluoromount-G Mounting Medium (Thermo Fisher Scientific). Image acquisition was performed on a Leica Thunder microscope, and images were processed with LasX software (Version 3.7.4.23463, Leica, Wetzlar, Germany) and Fiji.^38^ Cells were counted using the Cell Counter plugin and manually with Fiji.

### Quantitative RT-PCR

Cells were washed in PBS and dissociated into single cells with Accutase for counting. Cells were then pelleted and resuspended in PBS at a density of 1 to 10,000 cells per 5 μl. RNA extraction and cDNA synthesis were performed in a single step using the Super Script IV CellsDirect cDNA Synthesis Kit (Thermo Fisher Scientific) according to the manufacturer’s protocol. All qPCR reactions were performed in 384-well plates in 3 technical replicates with a 6 µl final reaction volume on a LightCycler 480 System II (Roche, Basel, Switzerland). Reaction mixes consisted of Power Track SYBR Green Master Mix 2X (Applied Biosystems, Thermo Fisher Scientific), primer pairs at 0.8 µMK concentration (**Table S2**), 2 µl of 1:5 diluted cDNA per reaction, and H_2_O to a final volume of 6 µl. The PCR programme consisted of an enzyme activation step at 95°C for 2 minutes, followed by 45 cycles of qPCR reaction at 95°C for 5 seconds and 65°C for 1 minute, and finally a melting curve from 60 to 97°C with 5 fluorescence acquisitions per °C. Expression levels were calculated by the comparative ΔΔCt method (2^−ΔΔCt^ formula), normalising to the Ct-value of the GAPDH housekeeping gene.

### Raman spectroscopy and imaging

Cells were washed in PBS and fixed in 4% PFA for 10 minutes at room temperature, followed by at least 3 further washes in PBS. Raman imaging was performed with a confocal Alpha-300R microscope from WITec GmbH (WITec Inc., Ulm, Germany). Excitation was assured by a frequency doubled Nd:YAG laser (Newport, Evry, France) at a wavelength of 532 nm. The incident laser beam was focused on to the sample through a 60× Nikon water immersion objective with a numerical aperture of 1.0 (Nikon, Tokyo, Japan). The acquisition time of a single spectrum was 0.5 s. The number of pixels in each image was 150 x 150.

Raman spectral data were initially acquired in the form of hyperspectral images. These datasets were subjected to K-means clustering to segment the images into major clusters representing distinct biochemical features of the cellular samples. The optimal number of clusters was determined by balancing cluster size and spectral representativeness. For each cluster, the mean spectrum was calculated and extracted as a representative spectrum. The collection of these mean spectra was considered to represent each sample for further analysis.^39^

The extracted spectra were subsequently processed using a fixed preprocessing pipeline (Le et al., 2026), including cosmic ray removal, baseline correction, optional smoothing, and normalisation. Baseline correction was performed using a peak stripping algorithm, while spectral normalisation was conducted using vector normalisation (L2 norm). Following preprocessing, Principal Component Analysis (PCA) was applied to explore spectral variability and identify key wavenumbers contributing to group discrimination. Based on these discriminative spectral markers, peak intensity ratios were calculated across all samples to monitor variations in cellular composition associated with these spectral features.

In this Raman data analysis study, we propose four major spectral regions: DNA-related, protein-related, lipid-related, and collagen-related regions. Each region is characterised by a score, defined as the mean value of selected Raman intensity ratios.

The DNA-related region is characterised by two prominent peaks at 727 cm⁻¹ and 793 cm⁻¹, corresponding to adenine and the O–P–O backbone vibrations of nucleic acids, respectively (**Table 1**). The DNA score is calculated based on ratios 727/1008 and 793/1008, where the intensities are normalised to the 1008 cm⁻¹ peak, a stable reference peak commonly used in Raman spectroscopy.

- The protein-related region is characterised by spectral features associated with protein vibrations (represented by 1008 and 1663 cm^-1^). These ratios are calculated relative to 2849 (CH₂ stretching), which is a stable and widely used reference peak in previous studies.^40^
- The lipid-related region is represented by three ratios 2849/1008, 2849/1663 and 2849/2945. The peak at 2849 cm⁻¹ (CH₂ stretching) is characteristic of lipids. These ratios are constructed to compare lipid-associated signals with strong protein-related peaks, such as phenylalanine (1008), amide I (1663) and CH_2_/CH_3_ stretching vibrations, thereby reflecting the relative contribution of lipid versus protein components.
- The collagen-related region is characterised by two ratios 920/2849 and 944/2849, associated with proline and hydroxyproline vibrations, which are characteristic amino acids of collagen (**Table 1**).

**Table 1.** Peak information in this study.^41,42^ These ratios were selected based on their contributions in the PCA loading profiles, with the aim of capturing biologically relevant variations while reducing dependence on absolute peak intensities.

| Practical Peaks | Interpretation |
| --- | --- |
| 727 | Adenine signal |
| 793 | O-P-O back bone of nucleic acid |
| 920 -944 | Proline and hydroxylproline of collagen |
| 1008 | Phenylalanine |
| 1454 | $\text{CH}_2$ stretching |
| 1680 | Amide I |
| 2849 | $\nu_s(\text{CH}_2)$, lipids, fatty acids |
| 2945 | $\nu_s(\text{CH}_2/\text{CH}_3)$, lipids, fatty acids, proteins |

## Supplementary Information

**Figure S1.**
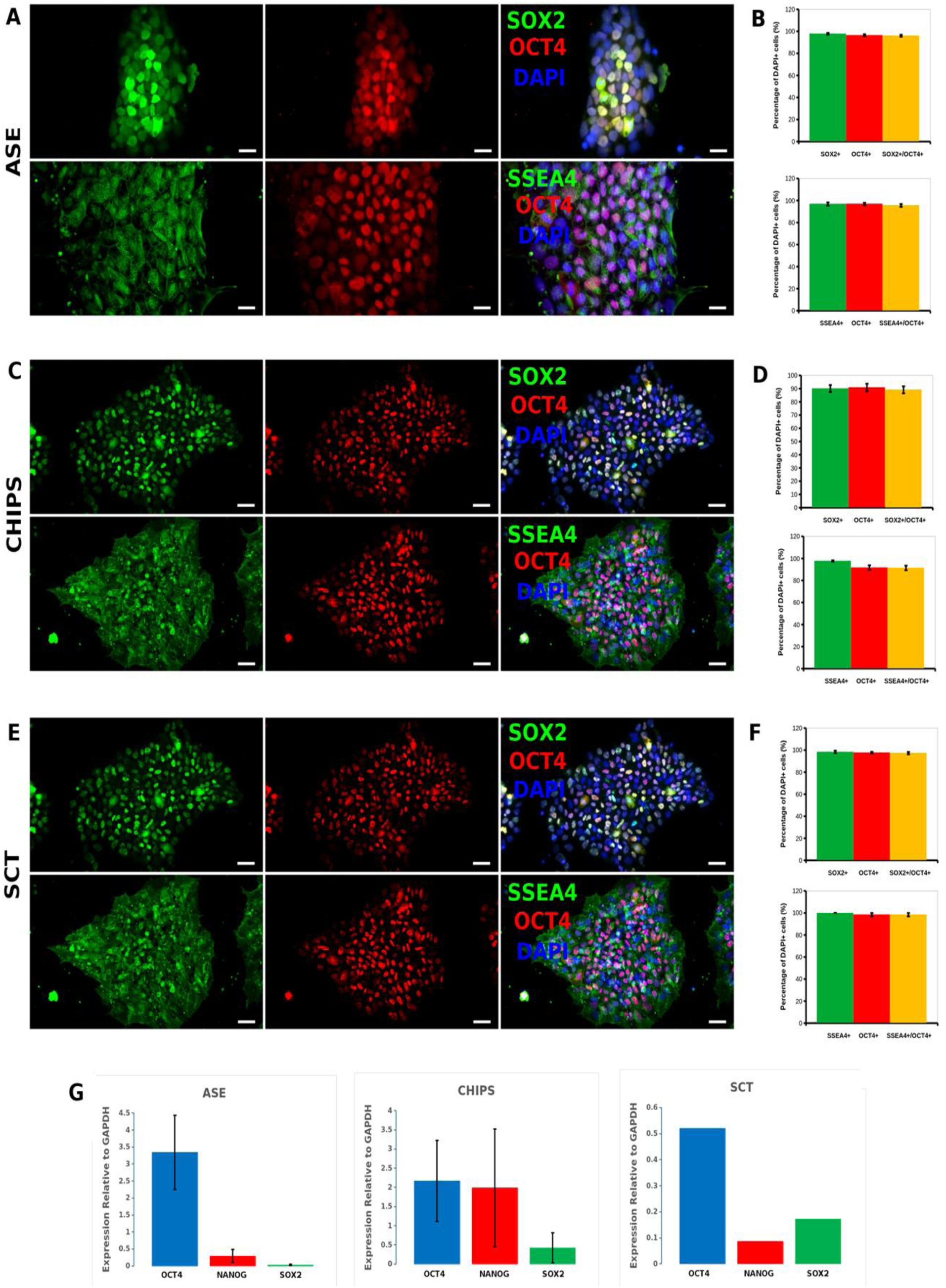
Immunofluorescence staining and quantification for SOX2, OCT4, and SSEA4 pluripotency markers for all cell lines used in the study (**A-F**). Scale bars = 20 µm. Quantification acquired from 3 - 4 images per line, normalised to DAPI and averaged, error bars = SEM. qPCR for *OCT4*, *NANOG*, and *SOX2* pluripotency markers normalised to *GAPDH* for all cell lines used in the study (**G**).

**Figure S2.**
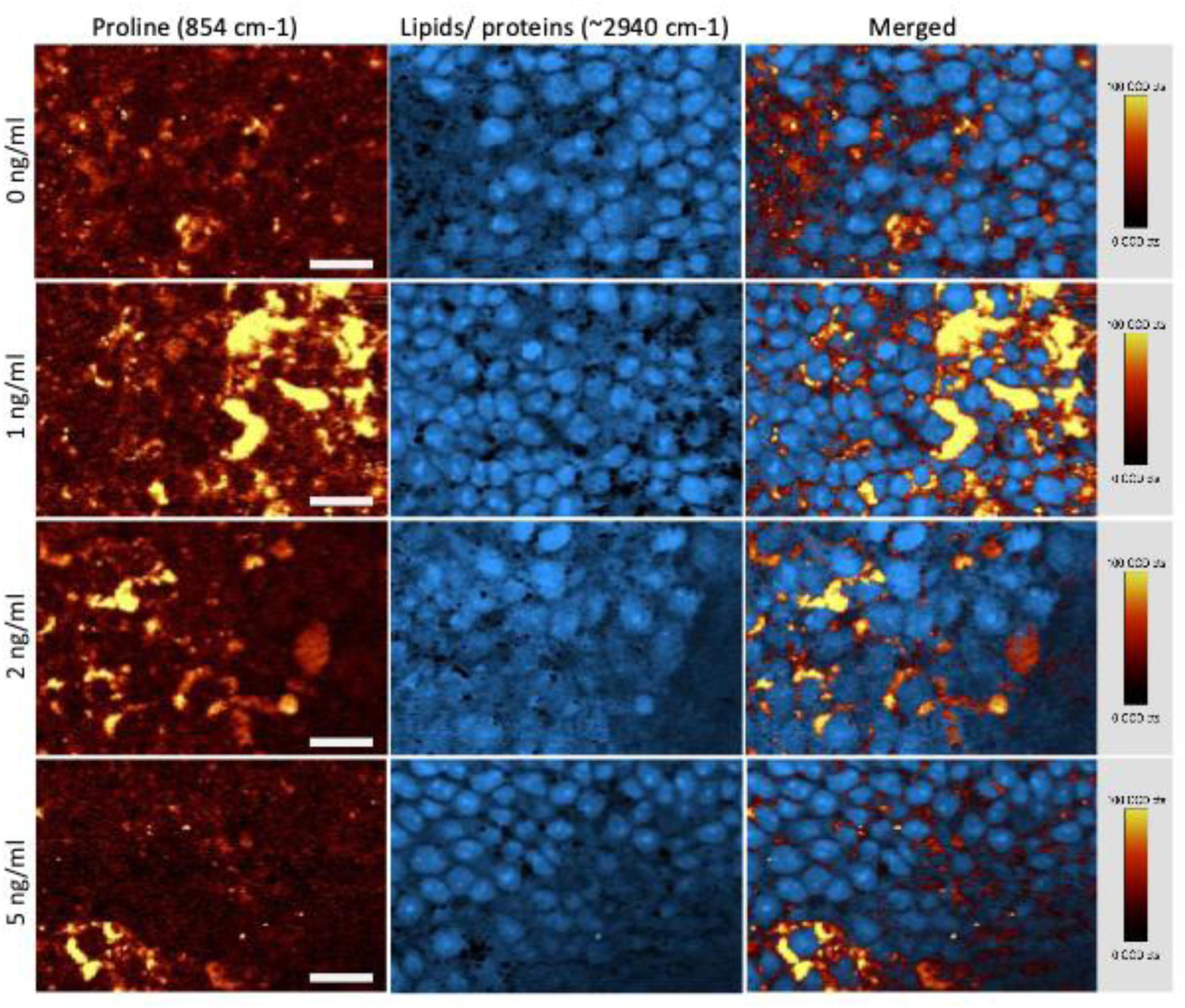
Monitoring iPSC differentiation toward placodal ectoderm derivatives under graded concentrations of BMP4 by Raman microscopy. False-colored images of the Proline peaks (854 cm-1), the C-H bond vibration corresponding mainly to lipids and proteins (∼2940 cm-1, not shown on the spectra), and the merging of both to show the different structures found by these two first images. Bars: 20 μm in all panels. The images are from cell differentiation of the ASE-9202 hiPSC line.

**Figure S3.**
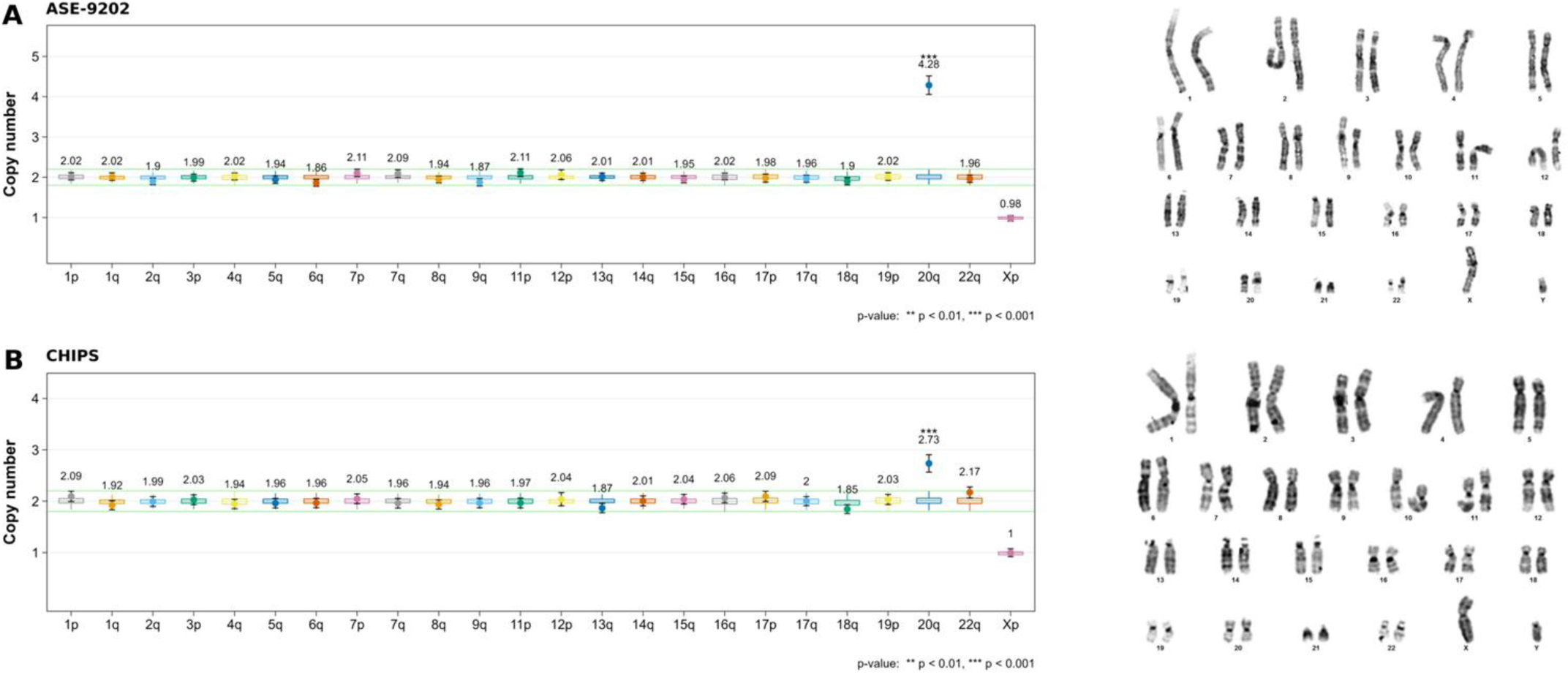
Copy number variation and cytoband results of the ASE (A) and CHIPS (B) cell lines.

**Table S1.**
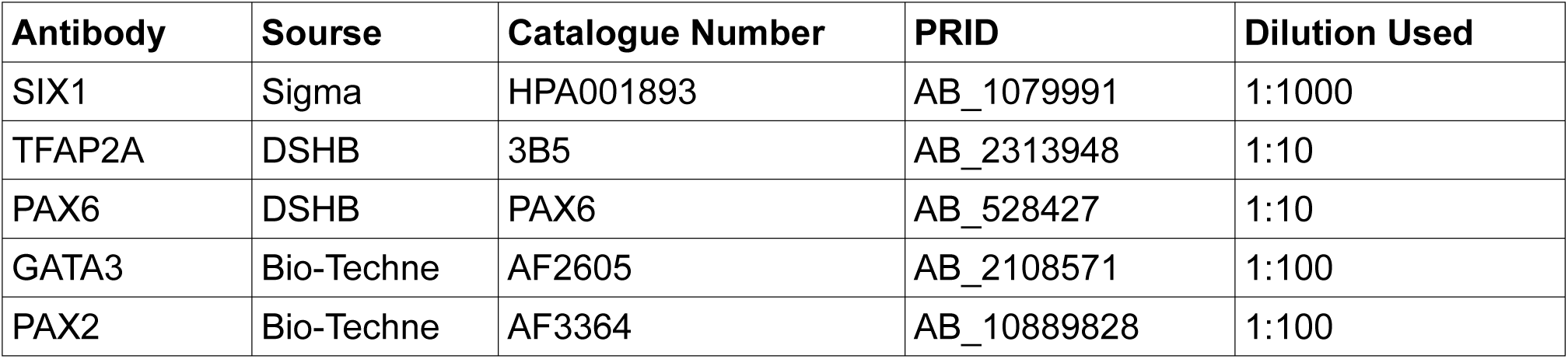
List of antibodies used in the study.

| Antibody | Source | Catalogue Number | PRID | Dilution Used |
| --- | --- | --- | --- | --- |
| SIX1 | Sigma | HPA001893 | AB_1079991 | 1:1000 |
| TFAP2A | DSHB | 3B5 | AB_2313948 | 1:10 |
| PAX6 | DSHB | PAX6 | AB_528427 | 1:10 |
| GATA3 | Bio-Techne | AF2605 | AB_2108571 | 1:100 |
| PAX2 | Bio-Techne | AF3364 | AB_10889828 | 1:100 |

**Table S2.**
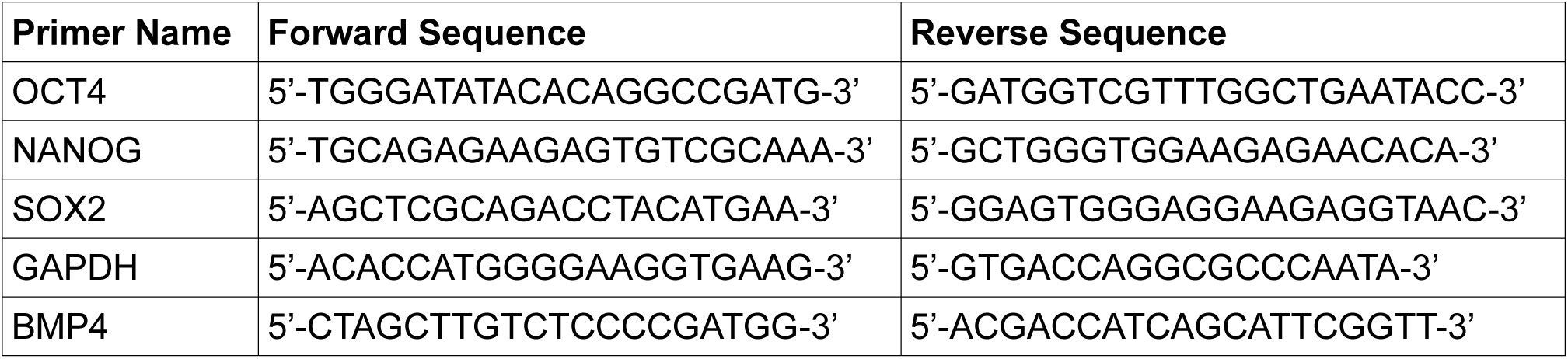
List of primers and sequences used in the study.

## Ethics approval and consent to participate

Not applicable.

## Consent for publication

Not applicable.

## Availability of data and materials

The datasets used and/or analysed during the current study are available from the corresponding author on request.

## Authors’ contributions

KC and AZ conceived and designed the experiments. KC, DV, LR, AD and EE performed the experiments. KC, DV and LR analysed and interpreted the data. KC, DV and AD drafted the manuscript. All authors read and approved the final manuscript.

## Acknowledgements

We thank the staff of the MRI-DBS-Optique facility (Elodie JUBLANC & Vicky DIAKOU) for the help with image acquisition and analysis and the RHEM-IGMM facility (Iria Gonzalez-Dopeso Reyes) for the help with histology.

